# Amphiphiles formed from synthetic DNA-nanomotifs mimic the step-wise dispersal of transcriptional clusters in the cell nucleus

**DOI:** 10.1101/2023.01.29.525851

**Authors:** Xenia Tschurikow, Aaron Gadzekpo, Mai P. Tran, Rakesh Chatterjee, Marcel Sobucki, Vasily Zaburdaev, Kerstin Göpfrich, Lennart Hilbert

## Abstract

Stem cells exhibit prominent clusters controlling the transcription of genes into RNA. These clusters form by a phase-separation mechanism, and their size and shape are controlled via an amphiphilic effect of transcribed genes. Here, we construct amphiphile-nanomotifs purely from DNA, and achieve similar size and shape control for phase-separated droplets formed from fully synthetic, self-interacting DNA-nanomotifs. Low amphiphile concentrations induce rounding of droplets, followed by splitting and, ultimately, full dispersal at higher concentrations. Super-resolution microscopy data obtained from zebrafish embryo stem cells reveal a comparable transition for transcriptional clusters with increasing transcription levels. Brownian dynamics and lattice simulations further confirm that addition of amphiphilic particles is sufficient to explain the observed changes in shape and size. Our work reproduces key aspects of the complex organization of transcription in biological cells using relatively simple, DNA sequence-programmable nanostructures, opening novel ways to control mesoscopic organization of synthetic nanomaterials.

**GRAPHICAL ABSTRACT:** 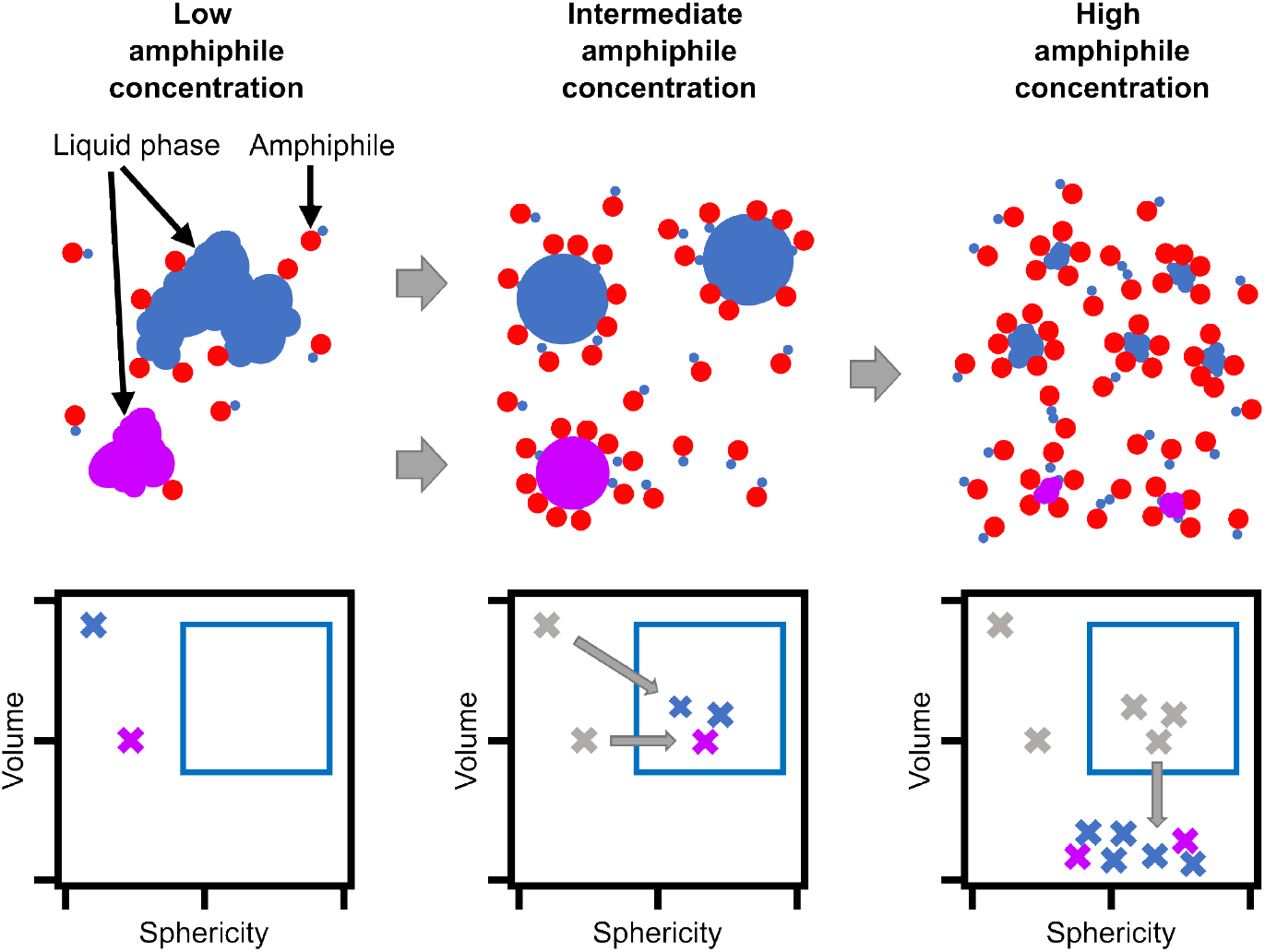

## Introduction

Biological cells need to compartmentalize their molecular components to ensure proper function. Conventionally, such compartmentalization is attributed to cellular organelles that are enclosed by membranes. More recently, processes based on liquid-liquid phase separation (LLPS) resulting from transient interactions were implicated in subcellular compartmentalization in biomolecular condensates.^1–3^ The transfer of biologically inspired compartmentalization via LLPS into biomimetic and synthetic systems yields remarkable design possibilities and performance increase.^4^ One example, a recently developed platform for fully programmable, multi-species phase separation is now available via DNA-nanomotifs, whose interactions are based on homology of “sticky ends” of exposed, self-complementary single-stranded DNA.^5, 6^ This platform has enabled the construction of a synthetic DNA segregation module that mimics chromosome separation during mitosis.^7^ Embedding into DNA-nanomotif condensates has also allowed the combinatorial detection of tumor biomarkers and up to 20-fold acceleration of DNA-based logic gates.^8, 9^

Condensates formed by canonical LLPS, commonly referred to as droplets, coarsen into increasingly larger droplets over time, whereas intra-cellular condensates are typically size-limited, or, in a state of microphase separation.^3, 10, 11^ Size-limited condensates can be established and maintained by different physical mechanisms, including active reaction processes,^12, 13^ targeted removal of aged condensates,^14^ or by provision of pinning centers at which condensates are localized.^15, 16^ Most commonly, however, microphase separation is attained by the addition of amphiphilic particles characterized by a dual affinity for two separating particle species.^17^ Size-control by amphiphiles can be seen in several biological example cases.^18–20^ A particularly well-studied case is the compartmentalization of transcription inside the cell nucleus. One prominent example is RNA polymerase II (Pol II), the enzyme complex responsible for transcribing most eukaryotic genes, which acts as a bivalent connecting particle between microphase-separated domains enriched in either DNA or RNA.^21–26^ An amphiphilic effect of ongoing transcription can also be seen by super-resolution microscopy of clusters of transcriptional machinery in stem cells.^27^ These clusters form by a phase-separation mechanism^28–35^ and unfold or even split into smaller clusters when they are engaged by genes that undergo transcriptional activation.^27, 36^ This effect can be reproduced in detail by a polymer model, in which transcribed genes are described as amphiphiles that associate with transcriptional clusters that, in turn, form by a liquid condensation mechanism.^27^ Here, we extend the synthetic DNA-nanomotif droplet platform by amphiphile-motifs and show that the effect of these amphiphile-motifs closely resembles the dispersal of embryonic transcriptional clusters.

## Results & Discussion

### Dispersal of RNA polymerase II clusters by an amphiphilic effect of transcribed genes

To allow for the comparison of the effect of DNA-based amphiphiles against a biological model system, we first characterize how transcriptional clusters are affected by the amphiphilic effects exerted by transcribed genes. In the zebrafish embryo, for a number of genes, transient visits to especially long-lived and prominent transcriptional clusters occur, which are tightly coupled with the transcriptional control of these genes by so-called “super-enhancers”.^27, 36^ As genes engage with transcriptional clusters and transcript elongation begins, transcriptional clusters unfold or even split into multiple parts (Figure 1A). This effect has been attributed to an amphiphilic property of genes that undergo activation in association with these prominent transcriptional clusters: while these genes exhibit an increased tendency to associate with transcriptional clusters, the beginning of transcript elongation additionally drives the exclusion of these genes from the transcriptional cluster.^36^ To obtain a fine-grained assessment of how increasing transcription levels relate to the dispersal of transcriptional clusters in zebrafish embryos, we labeled different regulatory states of Pol II by immunofluorescence. Recruited RNA polymerase II, which serves as a label for transcriptional clusters, can be detected via a specific phosphorylation mark of the RNA polymerase II C-terminal domain (Pol II S5P, Figure 1B). A different phosphorylation mark (Pol II S2P), can be used to detect currently elongating Pol II, which serves as an indicator of ongoing transcript elongation (Figure 1B). Fluorescence images recorded by instant-SIM super-resolution confocal microscopy^37^ revealed that the prominent Pol II S5P clusters were more unfolded and dispersed with higher levels of ongoing transcript elongation (Figure 1C). Assessing the fraction of large clusters at a given Pol II S2P level substantiates the impression that more large clusters occur at intermediate levels of transcript elongation, and clusters disperse at high levels of transcript elongation (Figure 1D). Quantifying the degree of unfolding via the sphericity of the Pol II S5P clusters also suggests the impression that higher Pol II S2P levels correlate with a less spherical shape of the Pol II S5P clusters (Figure 1C). Indeed, a comprehensive overview of Pol II S5P cluster shape was via sphericity-volume scatter plots revealed that a population of large (volume*≥* 0.08 *µ*m^3^) and round (sphericity*≥* 0.1) clusters is present specifically for intermediate levels of elongation activity (Figure 1E). At high levels of transcription, this population is reduced. Taken together, these analyses characterize how the amphiphilic effect of transcript elongation correlates with unfolding of transcriptional clusters, and, at the highest levels of transcript elongation, loss of large clusters.

**Figure 1:**
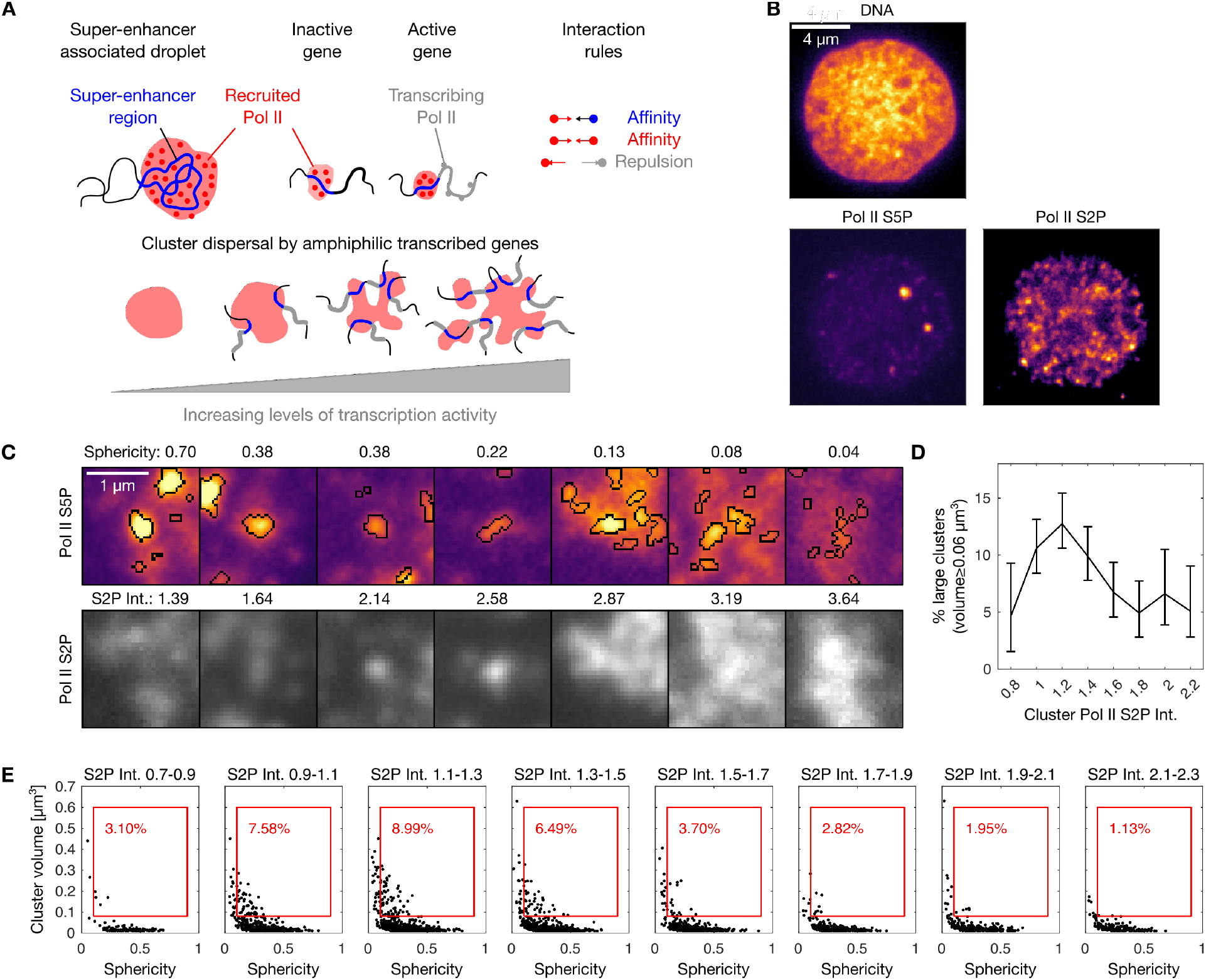
Transcribed genes act as amphiphile-like complexes that mediate the dispersal of RNA polymerase II clusters in nuclei of pluripotent zebrafish embryos. A)Illustration of the formation of clusters of recruited RNA polymerase II at super-enhancer regions of the genome and dispersal of clusters by an amphiphilic effect of transcribed genes. B) Representative three-color micrograph of an optical section through the middle of a cell nucleus in a pluripotent zebrafish embryo (sphere stage of development). Image data obtained by instant-SIM microscopy, color channels show DNA (Hoechst 33342), recruited RNA polymerase II (Pol II S5P, labeled by indirect immunofluorescence), and elongating RNA polymerase II (Pol II S2P, indirect immunofluorescence). C) Detail views of clusters of recruited RNA polymerase II, showing only the Pol II S5P and Pol II S2P channels. Segmentation was carried out in the Pol II S5P channel, as indicated in the top row. Sphericity was obtained from threshold-based segmentation of Pol II S5P clusters, S2P Int. refers to the mean intensity in the Pol II S2P channel, normalized by the median intensity of the nucleus that the cluster is contained in. D) Per-bin calculation of the percentage of large clusters in a given bin (volumes of 0.08 *µ*m^3^ or more, mean with 95% bootstrap confidence interval, data obtained from 99 nuclei pooled from 8 embryos). E) Segmented clusters were binned according to Pol II S2P intensity to generate sphericity-volume distributions. The percentage in red indicates binned clusters with volume ≥ 0.08 *µ*m^3^ and sphericity ≥ 0.1 (red box).

### Amphiphile-motifs formed purely from DNA can disperse droplets of self-interacting DNA-nanomotifs

After having characterized the dispersal of transcriptional clusters as a biological reference for the desired amphiphilic effects, we proceeded to develop amphiphiles that disperse droplets formed from DNA-nanomotifs. The previously developed phase-separating DNA-nanomotifs, called Y-motifs due their three-ended shape, self-interact via sequence-programmed sticky ends.^5^ Corresponding amphiphile-nanomotifs should also exhibit an affinity for the droplet-forming Y-motifs via sticky ends and, additionally, repulsion from the droplet phase (Figure 2A). Such a repulsion can be introduced, for example, by attaching negative charges via 5’-conjugation of a phosphate group, which resulted in a visually apparent reduction of droplet diameter, as is expected upon addition of an amphiphile (Figure 2B). An alternative approach to add negative charges for repulsion from the droplet phase, the extension by 102 thymines (poly-T extension), equally resulted in a visually apparent reduction of droplet diameter (Figure 2B). We theorized that the poly-T-based dispersal was due to a generic charge-based exclusion of the added DNA from the phase-separated droplets, which can only be overcome when nanomotif-nanomotif interactions, i.e., complementary base pairing, facilitate the retention within droplets. In line with this reasoning, the apparent dispersal of droplets by these amphiphile-motifs was abolished by the addition of 10 nt-long poly-adenine (poly-A) blocker oligomers, which can facilitate transient poly-T-poly-T cross-linking by complementary base pairing, similar to the sticky ends (Figure 2C). We used the poly-T-based amphiphile-motifs in the following steps of our study, as these amphiphile-motifs appeared to result in smaller droplets, and rely entirely on DNA-sequence effects, without chemical functionalization. To better understand the dispersal of Y-motif droplets, we added amphiphile-motifs to a suspension containing preformed Y-motif droplets, and visualized both species by time-lapse fluorescence microscopy (Figure 2D). Initially, amphiphile-motifs are homogeneously distributed throughout the volume not occupied by Y-motif droplets and excluded from Y-motif droplets (Figure 2E, 0 min inset). Amphiphile-motifs successively redistribute from outside to within the Y-motif droplets. Upon closer inspection, amphiphile-motifs exhibit a tendency to accumulate more strongly at the surface of Y-motif droplets relative to the inside of the Y-motif droplets, and a higher number of rounder droplets becomes apparent (Figure 2E, 30 min inset). Over time, the redistribution of amphiphile-motifs into the Y-motif droplets becomes yet more pronounced. Also, droplets are surrounded by a more clearly distinguishable layer of amphiphile-motifs and appear generally smaller, as would be expected after further splitting events (Figure 2E, 90 min inset). These observations suggest that amphiphile-motifs are successively redistributed into the Y-motif droplets and become incorporated into droplet surfaces. Over time, large droplets become increasingly energetically unfavourable and split into smaller droplets.

**Figure 2:**
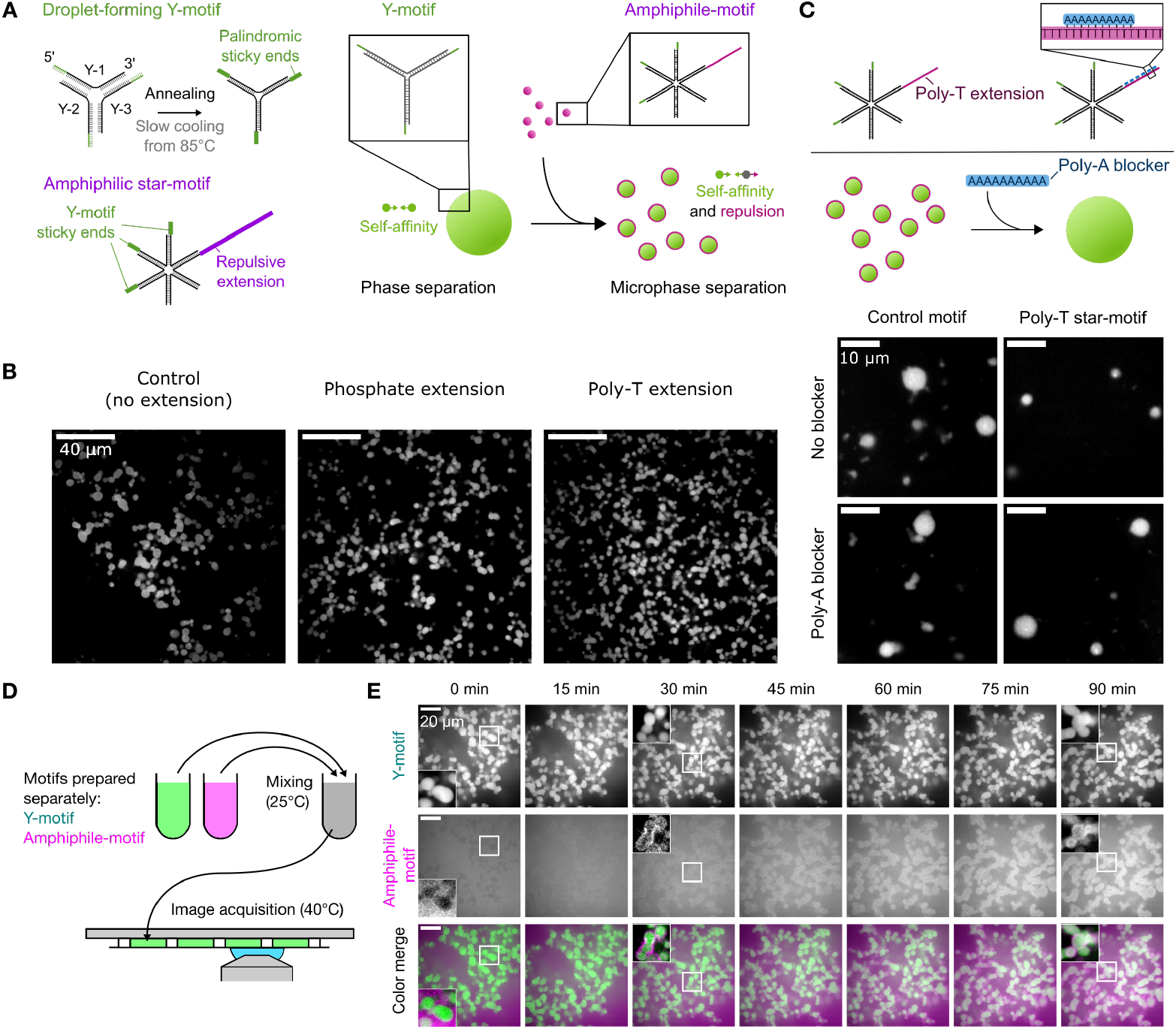
Amphiphiles for the dispersal of droplets formed from synthetic DNA-nanomotifs. A) Illustration of nanomotifs obtained by annealing of single DNA-oligomers, which consist of a 34 nt long core-structure, binding their neighboring strands, and a 8 nt long palindromic sticky end sequence. Amphiphile-motifs are generated by adding a repulsive extension to one vacant end of of a six-ended star motif. Y-motifs form phase-separated droplets due to the self-affinity established by sticky ends, which become dispersed upon the addition of amphiphile-motifs. B) Representative micrographs of Y-motif droplets in the presence of amphiphile-motifs. The negative charge is added via phosphate conjugation or the extension of one vacant end of the star-motif by 102 thymines (poly-T extension). Images are single confocal sections. C) Illustration of the charge-canceling effect from adding 10 nt long poly-A blockers. Micrographs show single confocal sections of Y-motif droplets in the presence of star-motifs without (control-motif) or with poly-T extension (amphiphile-motif), without or with addition of poly-A blockers. D) Y-motif droplets and amphiphile-motif solution were prepared in separate reaction tubes and subsequently mixed together for time-lapse imaging of the subsequent reorganization processes. E) Representative two-color time-lapse showing single confocal sections obtained from volumetric image stacks originally acquired with a 3-min time interval. Images 3D-deconvolved (Lucy-Richardson), Y-motif channel were corrected for photo-bleaching (histogram matching method), insets were contrast-adjusted to emphasize local details. Results were comparable over three independent repeats with two consecutive time-lapse recordings per repeat.

Our initial results suggest that amphiphile-motifs are recruited to Y-motif droplets before they exert their dispersing effect. In a time-course experiment with more precise temperature control (50*^◦^*C), indeed, Y-motif droplets initially took on rounder shapes, became increasingly smaller only over time until, ultimately, most droplets vanished (Figure 3A). The addition of control-motifs that lack the poly-T extension also induced rounding of Y-motif droplets, while, instead of becoming smaller, droplets continued to grow over time (Figure 3A).

**Figure 3:**
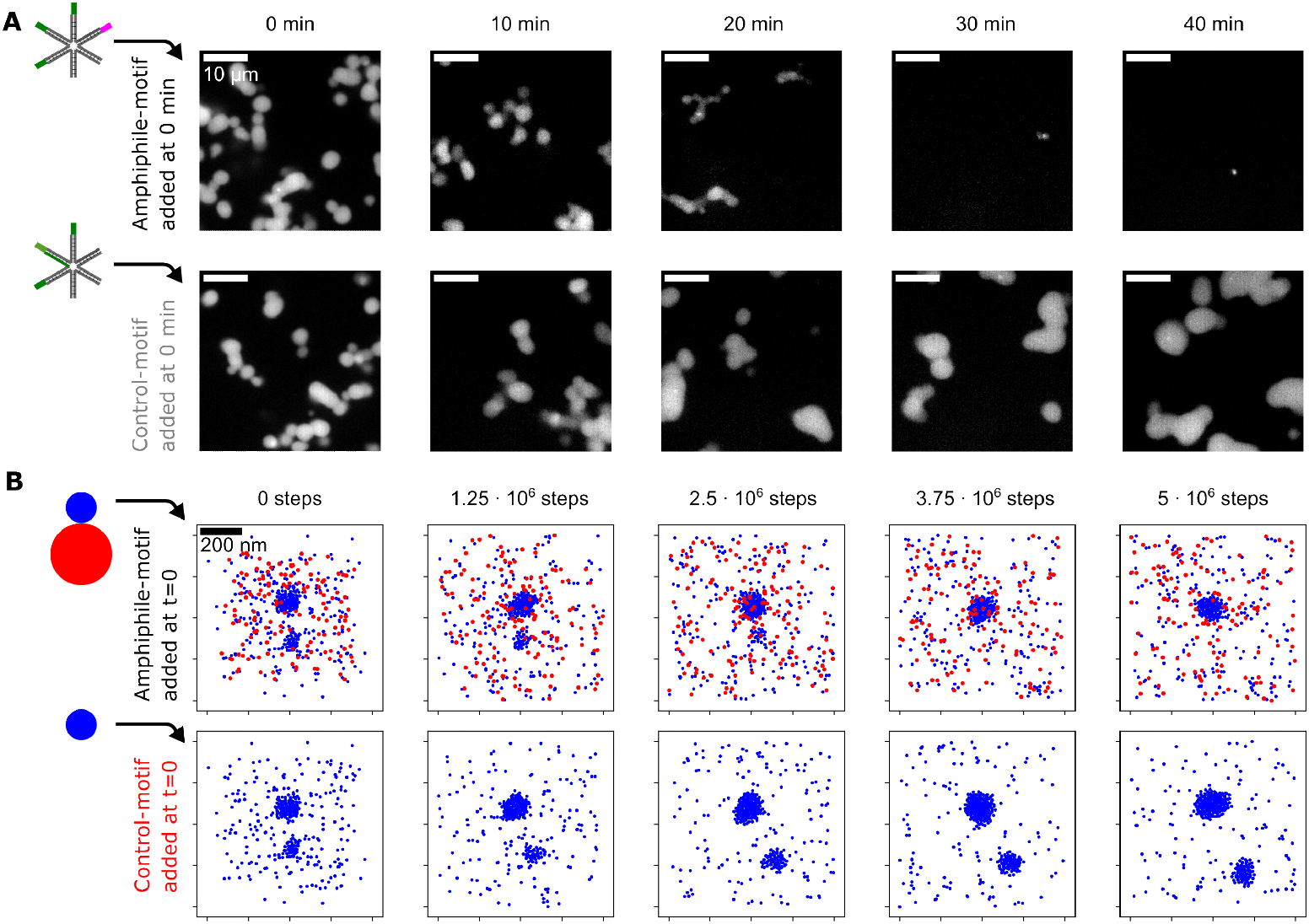
Time course of droplet dispersal upon addition of amphiphile-motifs. A) Representative confocal microscopy sections showing the distribution of Y-motifs in flow cells prepared at 10-min time intervals after amphiphile-motifs were added to preformed Y-motif droplets (0 min). Several tubes prepared in this manner were maintained at a stable temperature of 50*^◦^*C, control-motif samples were processed alongside amphiphilemotif experiments. B) To simulate the time-lapse experiment, either amphiphile-motifs or control-motifs were added to a simulation in a 1x1x1 *µ*m box containing preformed Y-motif droplets. Preformed droplets were generated by simulation with 450 Y-motifs for 10^6^ steps at 60*^◦^*C. 450 amphiphiles or control motifs (implemented as Y-motifs, displayed in blue) serving as control were then added and 5 10^6^ simulation steps at 40*^◦^*C were carried out. Images are 200 nm thick z-slices. Both simulations started from the same initial configuration.

We wanted to test whether attractive interactions between Y-motifs and amphiphile- motifs combined with a repulsive domain on the amphiphile-motifs are sufficient to explain these observations. We therefore implemented simulations based on Brownian dynamics, where Y-motifs were represented as spherical particles, which interact with other particles of the same kind via a harmonic well potential (Figure 3B, additional graphical summary see Figure S1). Amphiphile-motifs are represented as composite particles comprising one “Y-motif-particle” and one “tail particle”, which experiences harmonic repulsion from all other particles. The interaction potentials were adjusted so as to to recapitulate an experimentally observed transient increase in the number of droplets with increasing amphiphile-motif concentrations (Figure S2). We initialized a simulation with only Y-motifs, which we allowed to proceed until two differently sized droplets were formed, and subsequently added equal numbers of either amphiphile-motifs or control-motifs. The addition of amphiphile-motifs led to full dispersal of the smaller preformed droplet and prevented further growth of the larger droplet, whereas the addition of control-motifs resulted in continued growth for both droplets (Figure 3B). These observations already strongly indicated that, indeed, droplet dispersal can be attributed specifically to the repulsive poly-T tail. To consolidate these observations, we carried out a quantitative analysis of droplet number, volume, and sphericity for time-courses obtained experimentally and by simulation (Figure S3A-C and Figure S4A-C, respectively). In line with expectation, in both cases, addition of amphiphile-motifs to preformed Y-motif droplets induces a transient increase in volume and sphericity, followed by full droplet dispersal. Adding control-motifs also leads to an increase in droplet volume and sphericity, and, as expected, the subsequent volume reduction and dispersal do not occur.

### Addition of increasing amphiphile concentrations mimics the dispersal of RNA polymerase II clusters

Having verified that amphiphile-motifs induce the expected droplet dispersal, we investigated whether they could be used to mimic the dispersal of transcriptional clusters with increasing levels of transcript elongation in zebrafish embryos. To imitate increasing levels of transcript elongation, we prepared mixtures of Y-motifs with increasing concentrations of amphiphile-motifs (Figure 4A). In line with our previous observations, at all concentrations, amphiphile-motifs colocalized with Y-motif droplets and higher amphiphile concentrations were associated with smaller droplets (Figure 4B). In close agreement with observations made for Pol II clusters, also the droplet number transiently increases, then drops below the corresponding value without addition of amphiphile-motifs (Figure 4C). Also, the volume-sphericity distribution undergoes changes similar to Pol II clusters in zebrafish embryos for increasing levels of transcription (Figure 4D). In particular, the transient increase in the percentage of droplets with high volume and high sphericity mirrors the results for Pol II clusters. All observations were also visible in a second repeat of the titration experiment, indicating their reproducibility (Figure S5). Taken together, the changes in Y-motif droplets with increasing amphiphile-motif concentration resemble the changes in Pol II clusters in zebrafish embryos for increasing levels of transcript elongation.

**Figure 4:**
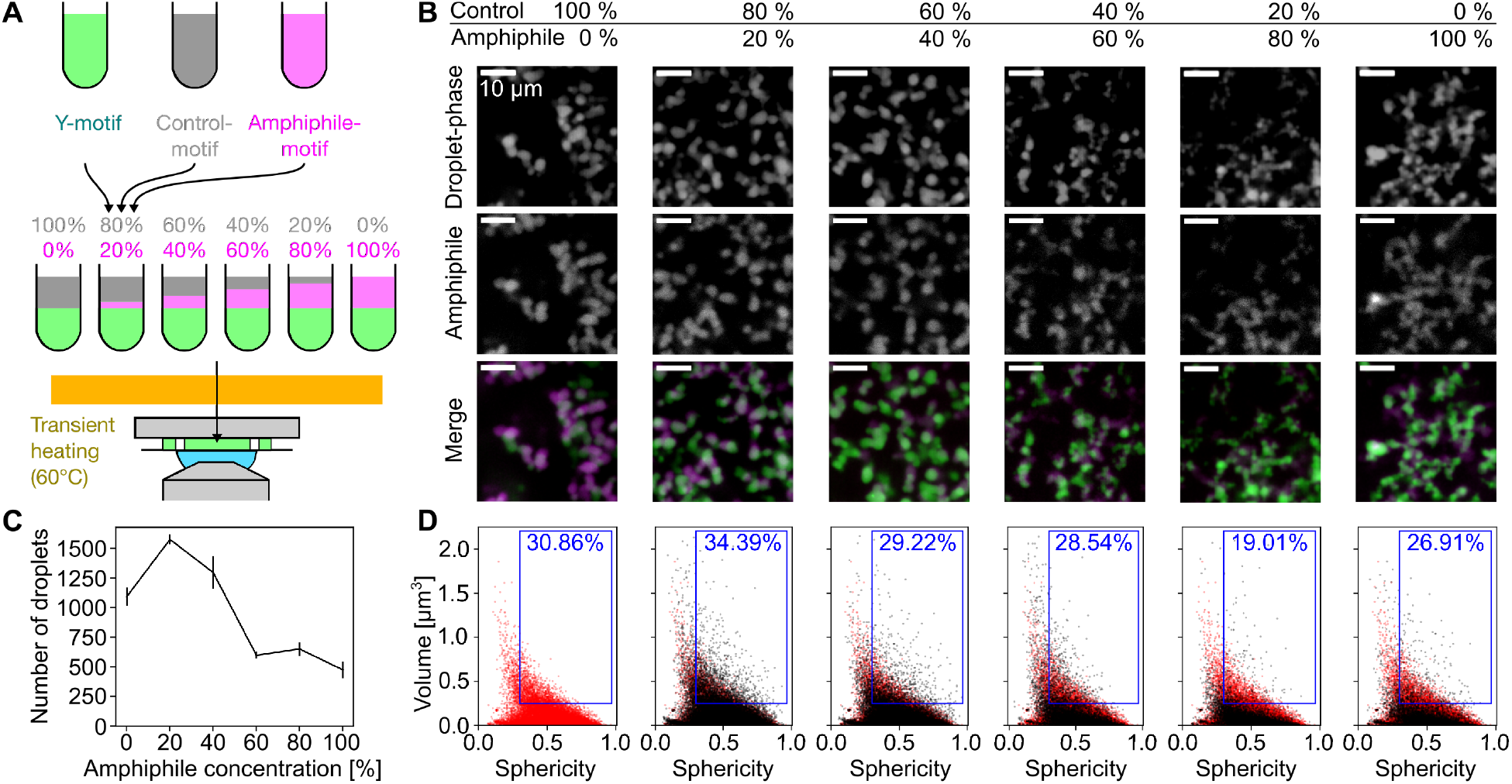
Increased amphiphile concentration leads to dispersal of phase-separated droplets. A) Y-motifs, control-motifs (no poly-T extension), and amphiphile-motifs were separately prepared, mixed at different percentages of amphiphile-motif, transiently heated (60*^◦^*C), and injected into flow cells for microscopy (ambient temperature 40*^◦^*C). B) Representative two-color micrographs of Y-motif and amphiphile-motif distributions at different amphiphile-motif concentrations. Micrographs are single confocal sections. C) Changes with amphiphile concentration of the number of droplets per field of view (mean±SEM, *n* = 14, 14, 13, 13, 14, 14 fields of view). D) Sphericity-volume distributions obtained from the analysis of confocal image stacks (image stacks were acquired at up to five fields of view per sample, n=15323, 22091, 18177, 7793, 8477, 6663 droplets, data pooled from *N* = 14, 14, 13, 13, 14, 14 total fields of view pooled from three independent experimental repeats). Small droplets (volume*≤* 0.125 *µ*m^3^) were excluded in calculation of percentages.

Finally, to assess whether our theoretical model can reproduce the dispersal of Y-motif droplets with increasing amphiphile-motif concentration, we simulated a titration experiment with increasing concentrations of amphiphile-motifs (Figure 5A). The volume fractions in the experiments with DNA-nanomotifs can be directly mapped to the particle number fractions in these simulations and the simulations are assigned physical length scales, so that results from simulations can be thoroughly compared to the experiments (Figure 5B). Visually, an increase in amphiphile-motif concentration leads to smaller Y-motif droplets and a more prominent accumulation of amphiphile-motifs on the surface of Y-motif droplets (Figure 5B). The number of detected droplets first rises and then drops, as seen in the experiments (Figure 5C). Sphericity-volume scatter plots show an increased fraction of large droplets with high sphericity, followed by a decrease of this fraction for even higher amphiphile-motif concentrations (Figure 5D). Taken together, our theoretical model largely captures the experimental observations for increasing concentrations of amphiphile-motifs, and, by extension, for transcriptional clusters with increasing levels of transcription.

**Figure 5:**
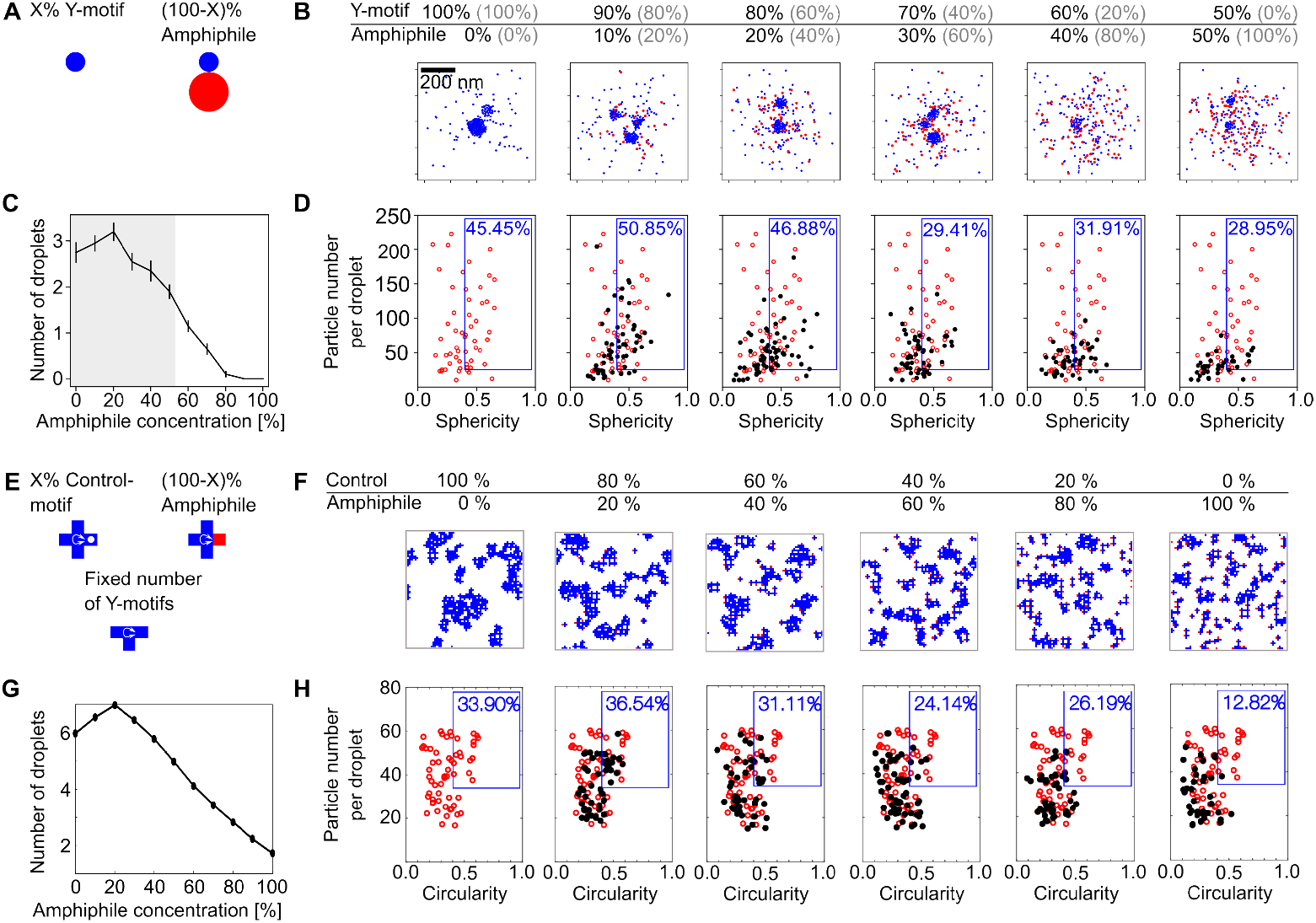
Amphiphile titration in interacting-particle and lattice simulations. A) To-scale drawing of Y-motifs and amphiphile-motifs used in the interacting-particle simuations. B) Particle-based simulations with 450 particles in total were used to represent the titration experiment. The fraction of amphiphile was increased from 0% to 100% in steps of 10%, the remaining fraction comprising of Y-motifs. Initially, all particles were distributed in a spherical volume of 200 nm inside a simulation box of 1x1x1 *µ*m. 10^6^ simulation steps were then carried out and the final configuration was evaluated. Due to the different methods of assigning percentages (number fractions in the simulation, volume fractions in the experiment), we provide corresponding percentages from the experiment in grey. One example of a z-slice (thickness 200 nm) of the final configuration is shown for each case. C) Number of droplets per simulation, grey regions correspond to the range investigated in the experiment, mean SEM (*n* = 20 simulations). D) Distributions of sphericity against particle number per droplet from 20 simulations. The blue box contains a percentage of droplets with sphericity higher than 0.4 and more than 25 particles per droplet. E) Lattice-based simulations with three different particles types were used to represent the titration experiment. T-shaped and cross-shaped DNA motifs perform diffusion and undergo stochastic rotation. Amphiphiles have one repulsive interacting arm (red), which is replaced by a neutral arm (white dot) for controls. F) Images show representative configurations of simulations carried out on 64 64 two-dimensional lattice. G) Number of droplets per simulation, mean SEM (*n* = 25 simulations). H) Distributions of circularity against particle number per droplet from 25 simulations. The percentage is for droplets with circularity higher than 0.40 and more than 35 particles per droplet.

To test the generality and robustness of our theoretical results, we implemented phenomenological lattice-based simulations including self-interacting particles and amphiphilic particles, thereby focusing on essential interaction rules (Figure 5E). The simulation is accomplished on a 2-dimensional square lattice with different types of particles representing the different types nanomotifs. Particles could not overlap and moved according to a dynamic Monte-Carlo algorithm. Different affinities were assigned to the tips of the particles to represent the different nanomotifs. We consider three types of particles: three-armed particles with self-affinity analogous to the DNA Y-motifs; amphiphilic particles with an additional, repulsive arm; and control particles similar to the amphiphiles, except that the additional arm is neutral (Figure 5E). Affinities and the particle density were adjusted so that self-interacting particles behave as a phase-separating liquid.^38–40^ When we simulate the titration of amphiphile particles, large phase-separated domains are dispersed as the fraction of amphiphile particles is increased, as seen in the interacting-particle simulations (Figure 5F). Further, a transient increase in droplet number and in the fraction of large, round droplets is observed, in line with the interacting-particle simulations and experiments (Figure 5G,H). This agreement between lattice simulation and interacting-particle simulation results further supports that, indeed, the dispersal of phase-separated droplets by increasing amphiphile concentrations can be attributed to the same essential set of interaction rules. The robustness of our theoretical results is further supported by a quantitative analysis of number of droplets, particle number per droplet, and circularity for time-lapse simulations, which matches experimental and interacting-particle simulation results (Figure S6).

## Conclusion

Taken together, in all experimental and theoretical systems examined in this study, a stereotypical dispersal process was observed with increasing amphiphile concentrations. An increase in the fraction of large, round droplets and the total number of droplets can be observed at low amphiphile concentrations. At sufficiently high amphiphile concentrations, splitting of droplets leads to a dispersal beyond the detection limit. In our understanding, this stereotypical behavior can be traced back to the similarity in the underlying amphiphile-mediated dispersal process. For low amphiphile levels, droplets grow as amphiphile particles are incorporated. Round, convex shapes become energetically favourable, as they maximise the distance between the repulsive, surface-attached parts of the amphiphile.^41^ For increasing amphiphile concentrations, the requirement for increased droplet surface area to accomodate the additional amphiphile particles leads to splitting and, ultimately, dispersal of droplets.^17^ In summary, our results show how an amphiphilic DNA-nanomotif constructed solely on the basis of sequence interactions, without chemical functionalization, can be used to control the size of phase-separated droplet compartments or even induce complete dispersal of these compartments, similar to what is observed for transcriptional clusters in stem cells. Several additional features not reflected in the synthetic DNA-motifs used in our study could be considered in future to achieve more realistic synthetic reimplementations of transcriptional clusters. Transcription in eukaryotes proceeds via several adjacent compartments at the scale of a few 100 nm, which harbor the consecutive steps of transcription control, transcript elongation, and splicing of the resulting transcripts.^42–46^ The ability to implement several, orthogonally interacting DNA-motifs based on different sticky end sequences allows a similar formation of distinct droplet species, whose fusion and separation can be controlled by additional connector motifs.^5, 7^ The formation of transcriptional clusters in pluripotent cells involves super-enhancers as condensation surfaces,^27–31, 43, 47–51^ whereas the Y-motif droplets in our study form without such condensation surfaces. Such surfaces could be provided by long DNA strands produced with the rolling circle amplification (RCA) method, which was previously used to generate DNA-only liquid droplets.^52^ Lastly, the amphiphilic DNA-nanomotifs in our synthetic system are catalytically inactive, while the ongoing synthesis and transport of RNA can distinctly change the stability of transcriptional clusters^27, 53^ as well as the mesoscopic organization of different steps of the transcription process and chromatin.^21, 54, 55^ In conclusion, our work illustrates how principles of intracellular compartmentalization can be translated into novel, biomimetic concepts for the control of synthetic biological systems and materials.

## Supporting information

OligomerSequences

## Acknowledgements

X.T., A.G., M.S., and L.H. are supported by the Helmholtz Program Natural, Artificial, and Cognitive Information Processing (NACIP). X.T. and L.H. received seed funding from the KIT Center Materials for Technology and Life Sciences (MaTeLiS). M.P.T and K.G received funding by the Hector Fellow Academy, the Federal Ministry of Education and Research (BMBF), and the Ministry of Science Baden-Württemberg within the framework of the Excellence Strategy of the Federal, and State Governments of Germany and the Max Planck Society. R.C. and V.Z. were supported by the ”Life?” initiative of the Volkswagen Stiftung.

## Supporting Information Available Methods and Materials

### Zebrafish husbandry and collection of embryos

All zebrafish husbandry was performed in accordance with the EU directive 2010/63/EU and German animal protection standards (Tierschutzgesetz §11, Abs. 1, No. 1) and is under supervision of the government of Baden-Württemberg, Regierungspräasidium Karlsruhe, Germany (Aktenzeichen35-9185.64/BH KIT).

### Immunofluorescence staining of zebrafish embryo animal caps

Zebrafish embryos in the sphere stage were collected as previously explained.^27^ In particular, wild-type adult fish (AB strain from the Zebrafish International Resource Center, maintained at the European Zebrafish Resource Center) were set up for spontaneous mating, eggs were collected and chorions removed by treatment with pronase. Embryos were maintained in 0.3X Danieau’s solution, and transferred into 0.3X Danieau’s supplemented with 2% formaldehyde and 0.2% Tween-20 for fixation. Permeabilization was carried out by 15 min exposure to 0.5% Triton X-100 in phosphate-buffered saline (PBS, Dulbeccos’s formulation) at room temperature. After two washes in PBS with 0.1% Tween-20 (PBST), 30 min of blocking with 4% Bovine Serume Albumin (BSA) in PBST followed. Samples were then incubated at 4*^◦^*C overnight with primary antibodies against Pol II S2P (Abcam ab193468, rabbit

IgG monoclonal, clone EPR18855, dilution 1:300) and Pol II S5P (Active Motif, rat IgG monoclonal, clone 3E8, dilution 1:300) in 4% BSA-PBST. After 3 washes in PBST, the samples were again incubated overnight with secondary antibodies (ThermoFisher, anti-rat polyclonal conjugated with Alexa 594, dilution 1:300; ThermoFisher anti-rabbit polyclonal conjugated with STAR RED, dilution 1:300) in 4% BSA-PBST. Samples were washed 3 times in PBST, post-fixed in 2% formaldehyde in PBS for 15 min at room temperature, again washed 3 times in PBST. Mounting was carried out by placing animal caps immersed in non-setting mounting media (VectaShield H-1000) supplemented with 1.6 *µ*M Hoechst 33342 between adhesive strips spaced 2-4 mm apart, which were covered with thickness-selected #1.5 glass cover-slips. The assembled sample was then sealed with nail polish.

### Preparation of flow cells on microscope slides

Flow cells were prepared by placing strips of double-sided tape on glass microscope slides to form several channels. To block surfaces for binding, the flow cells were incubated with 2% bovine serum albumin (BSA) for 5 min. After removing the remaining solution with a chemical wipe, the microscope slides were air-dried. Then cover slips were placed on the microscope slides. For imaging, the flow cell channels were filled with sample solution and sealed with silicone grease.

### Preparation of DNA-nanomotif suspensions

#### Oligonucleotide preparation

Oligonucleotides were purchased from biomers.net and Integrated DNA Technologies (IDT) with purification by standard desalting (unmodified strands) or HPLC (fluorophore-tagged strands). The individual strand sequences were taken from previous work^5^ and adapted as detailed in Supplementary Table S1. Individual oligonucleotide preparations were dissolved in 1X TE at 200 *µ*M and stored at −20°C up until preparation of mixtures needed for a given experiment.

#### Nanomotif formation

To produce the different nanomotifs (Figure 2A, B), the corresponding DNA-oligomers were separately mixed together in 25 *µ*l of 2X TE with 700 mM NaCl. The solutions were then heated up with a thermocycler (Eppendorf Mastercycler X50) to 85°C for 3 min, before decreasing the temperature at a rate of −1°C/min to 40°C. How the different star-motifs affect the Y-motif droplets was investigated by first preparing all components separately (5.0 *µ*M Y-motif and 1.65 *µ*M star-motif), then mixing both parts in equal proportions and heating up to 60°C before decreasing the temperature at a rate of −1°C/min. Annealed nanomotifs were used immediately.

#### Inhibition of amphiphile with poly-A blocker elements

Nanomotifs were prepared separately, (10.0 *µ*M Y-motif and 5.0 *µ*M amphiphile-motif). Then, 4.44 *µ*M Y-motif and 2.22 *µ*M amphiphile-motif were mixed together with 22.2 *µ*M of a 10-nt poly-A blocker (for sequences see Supplementary Table S1). Control solutions were prepared by the substitution of the poly-A blocker solution with an equivalent amount of water. The samples were heated up to 60°C, then the temperature was decreased at a rate of −1°C/min to 40°C before imaging the samples.

#### Time-lapse of amphiphile-droplet interaction

Time-lapse experiments were performed by first preparing 20.0 *µ*M Y-motif and 15.0 *µ*M of amphiphile-motif in separate tubes, then mixing 10.0 *µ*M of the Y-motif solution with 7.5 *µ*M of the amphiphile-motif solution (concentrations in final mixture). The mixture was then added to preheated (40°C) flow cells and imaged at 40°C for up to 4 h.

#### Time-course of amphiphile-droplet interaction with incubation in a thermocycler

The time-lapse experiments were repeated at 50°C, assuming that the nanomotifs reorganize faster at elevated temperature. Here, 10.0 *µ*M of Y-motif and 15.0 *µ*M of amphiphile-motif and control-motif were prepared separately, before mixing 5.0 *µ*M Y-motif and 7.5 *µ*M amphiphile-motif or control-motif together (concentrations in final mixture). Several tubes were prepared and incubated at 50°C in a thermocycler. Every 10 min, an amphiphile-motif and a control-motif sample were retrieved from the thermocycler and injected into preheated flow cells (40°C) for imaging (ambient temperature 40°C).

#### Titration of amphiphile-motifs

Titration experiments were conducted by preparing 30.0 *µ*M of Y-motifs and 22.5 *µ*M of amphiphile-motifs and control-motifs in 3 separate tubes. Then, 6 tubes of 15 *µ*M Y-motif and 11.25 *µ*M amphiphile and control-motif were mixed together. In the first tube, only 11.25 *µ*M of control-motif were added. In the following tubes, the control-motif was replaced with amphiphile-motifs in 20%-steps, so that the last tube only contained amphiphile-motifs and Y-motifs. These tubes were then heated up to 60°C and cooled down at a rate of −1°C/min to 40°C. For imaging, the different samples were injected in prewarmed flow cells (40°C) and imaged at 40°C.

#### Imaging procedure

Samples were visualized using an instant structured illumination microscope (iSIM) from VisiTech. Before injection, flow cells were heated up to 40°C. Microscopy of the samples was carried out in a heated microscope stage enclosure at a temperature of 40°C.

### Microscopy and image analysis

#### Zebrafish embryos

Confocal microscopy images were obtained using a VisiTech iSIM microscope, which is a commercial implementation of a microscope based on the the instant-SIM principle.^37^ The microscope is based on a Nikon Eclipse Ti2 microsope body. A motorized correction collar oil immersion objective (CFI SR HP Apo TIRF 100XAC Oil, NA 1.49) was used. Images were acquired with a single ORCA-Flash4.0 V3 camera with sequential acquisition of color channels for maximal positional accuracy and minimal cross-talk. To limit the analysis to conventional gene and enhancer loci, nuclei with visible miR-430-associated foci were excluded from the data set.^56–58^

The analysis of fluorescence intensities and shapes of Pol II clusters was carried out following our previous work,^27^ using MatLab for analysis and the Open Microscopy Environment bioformats plugin for data import.^59^ In brief, a two-step segmentation pipeline was applied, first segmenting cell nuclei by Otsu-thresholding of the DNA-channel, followed by robust background threshold segmentation of Pol II clusters from the Pol II Figure channel in each detected nucleus. For each detected cluster, the geometric characteristics volume and sphericity and the mean intensities in the Pol II S5P and the Pol II S2P channel were calculated under consideration of the 3D-shape segmentation mask. Mean intensities were normalized against the median intensity of the containing nucleus to compensate for variability in sample staining and image acquisition between different nuclei and samples. Sphericity was calculated as

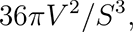

where *V* is the volume and *S* the surface area of a given 3D object

The raw data and the image analysis code for Figure 1 is deposited on Zenodo: https://doi.org/10.5281/zenodo.5242771

#### Nanomotif suspensions

Nanomotifs were imaged using a water immersion objective (Apo LWD 40x WI *λ*S DIC N2, NA 1.15). Images were recorded using dual ORCA-Flash4.0 V3 cameras with simultaneous two-channel acquisition to avoid displacement of objects due to consecutive acqusition of color channels.

The images were further processed with the software Fiji.^60^ Raw images containing the droplets formed by Y-motifs were resliced in the z-axis to generate cubic voxels and then filtered using Gaussian Blur 3D with the parameters *σ* = 2,2,2 pixels for x, y, and z, respectively. To obtain image data for the volume, sphericity, and elongation of the detected droplets, the plugin MorphoLibJ was used.^61^ Images were processed with the ”Morphological Segmentation” tool as object images with the gradient type “Morphological”, gradient radius = 2 and a tolerance = 100. Images were displayed in the “Catchment basins” format. The background was removed by using the command “Remove Largest Object”. Images were further analyzed using the tools “Analyse Regions 3D” for the volume and sphericity (surface area method = Crofton (13 dirs.), Euler connectivity = 6).

The raw data and the image analysis code is deposited on Zenodo.

Comparison of different amphiphiles (Figure 2B): https://doi.org/10.5281/zenodo.7572656

Blocking of poly-T amphiphiles (Figure 2C): https://doi.org/10.5281/zenodo.7572624 Continuous time-lapse on the microscope, 40*^◦^* (Figure 2E): https://doi.org/10.5281/zenodo.7572182

Manual time-lapse, 50*^◦^* (Figure 3A and Figure S3A-C): https://doi.org/10.5281/zenodo.7572230

Titration poly-T amphiphile (Figure 4B-D and Figure S5A, B) https://doi.org/10.5281/zenodo.7573452

### Interacting-particle model

#### Interacting-particle dynamics in ReaDDy

Brownian dynamics is a common approach to describe particles subject to both deterministic forces in a Newtonian sense and stochastic forces that result in Brownian motion. Particle motion is determined by an overdamped version of a stochastic differential equation called Langevin equation:

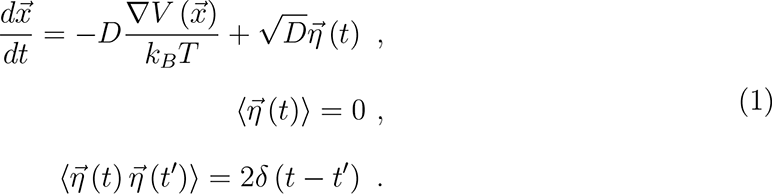

The particle position is denoted by 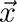, the diffusion constant by *D*, the Boltzmann constant times temperature by *k_B_T* and a delta-correlated Gaussian noise term by 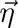. Interactions are summed up in the potential *V* . The fluctuation-dissipation theorem (*σ*^2^ = *ξk_B_T* ) and specifically the Einstein relation 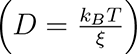 were applied to relate the friction coefficient *ξ*, the noise amplitude *σ*, and the diffusion constant. The noise contribution is drawn from a Gaussian with zero mean and variance of one 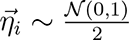 .

ReaDDy (short for Reaction Diffusion Dynamics) is a software package capable of numerically integrating the Langevin equation (1) for individual particles. Importantly, it also allows the user to include various potentials for interactions between the particles and chemical reactions that can convert, create, or delete particles. This kind of simulation is often referred to as interacting-particle reaction dynamics (iPRD) and has proven versatile enough to model and simulate a variety of biochemical systems. ReaDDy, in version 2.0.12, was used for all simulations based on Brownian dynamics. A more in-depth introduction to ReaDDy can be found in.^62^

The Euler-Maruyama method is used to integrate the Langevin equation. The evolution from a position 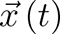 to 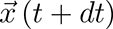 can be written as:

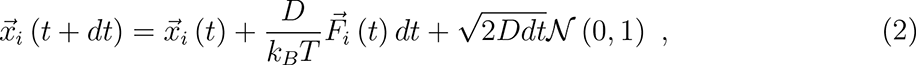

where the effects of external forces and interaction forces with other particles are summed up in 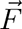 :

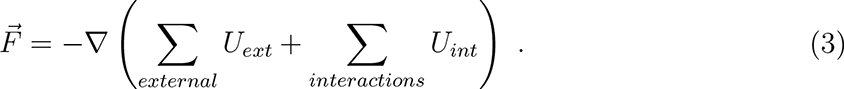

A variety of potentials can be used to model inclusion or exclusion effects and attraction or repulsion between particles. Examples include Lennard-Jones or harmonic pair potentials, spherical barriers or inclusion potentials. A detailed summary can be found on the website of ReaDDy (https://readdy.github.io/system.html#potentials).

ReaDDy is accessible via a Python interface, however, the core functionality is written in C++. This way, the user benefits from Python’s accessibility without sacrificing optimisation for speed.

#### Parameters of simulation

The droplet simulation is a coarse-scale, effective model of the DNA-nanomotif system. Our aim was to find a parameter space for particle interaction that reproduces the experimental results. Particle concentrations and sizes are thus not directly comparable to the DNA-system.

Our model for the Y-motif is a spherical particle with diffusion constant *D_y_* = 0.0352 nm^2^*/*ns obtained from the Einstein relation. Interaction with other Y-motifs is defined by a harmonic well potential:

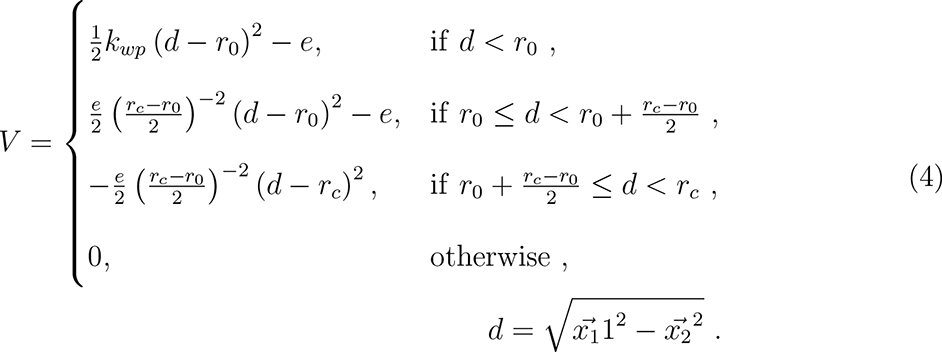

After testing various parameter settings, we decided to use the following for our simulations:

- spring constant of repulsive part: *k_wp_* = 10^−20^ J*/*nm^2^
- depth of well: *e* = 2 *· k_B_T*
- position of minimum: *r*_0_ = 20 nm
- cut-off distance: *r_c_* = 30 nm

The amphiphile particle comprises of one particle identical to the Y-motif model and one ”tail particle” that is repulsed by all other particles via a harmonic potential:

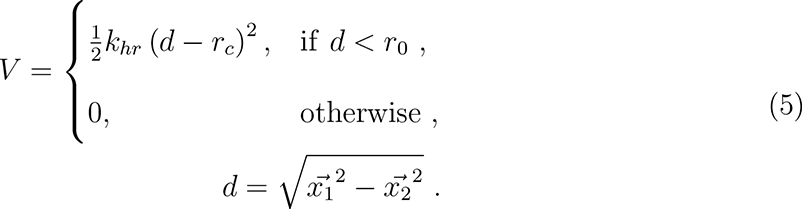

The following parameters were used:

- spring constant: *k_hr_* = 10^−20^ J*/*nm^2^
- cut-off distance: *r_c_* = 30 nm

The diffusion constant of the ”tail particle” was set to *D_y_* = 0.0175 nm^2^*/*ns to model a larger effective volume.

A bond potential defines the interaction between the two sub-particles of our amphiphile model:

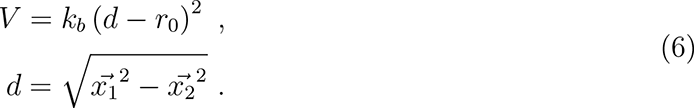

Here, we used the following parameters:

- spring constant: *k_hr_* = 10*^−^*^20^ J*/*nm^2^
- minimum distance: *r*_0_ = 31 nm

All particles are also subject to an inclusion potential defined by:

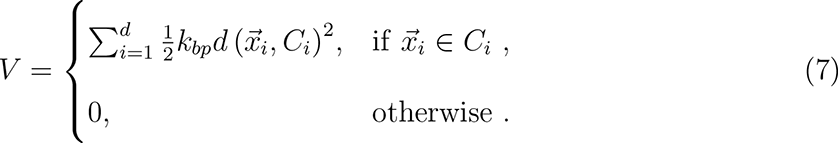

The term 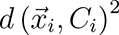 represents the shortest distance between the particle position *x7_i_* and the boundary of a cube *C_i_* = [*origin_i_, origin_i_* + *extent_i_*].

We used the following parameters:

- spring constant: *k_bp_* = 10*^−^*^20^ J*/*nm^2^
- extent of box = 800 nm

The box was centered in a simulation volume of extension 1x1x1 *µ*m.

#### Simulation of time series

Simulations for the time series comprised of two steps: In the first step, 450 Y-motif particles were randomly placed in a spherical volume of radius 200 nm and simulated for 10^6^ time steps of 0.5 ns and 60*^◦^*C. The final configuration containing preformed droplets was used for the second step. Here, 450 additional Y-motifs (as control case) or amphiphile-motifs were randomly added into a volume of 0.8x0.8x0.8 *µ*m, without creating particle overlaps. The system was then simulated for 5 *·* 10^6^ steps of 0.5 ns and a reduced temperature of 40*^◦^*C. This process was repeated for 10 independent simulations, using the standard interaction parameters defined above.

#### Simulation of titration

Simulations for the titration split 450 particles between the models for Y-motif and amphiphile-motif. The amphiphile percentage was increased from 0% to 100% in steps of 10%. 20 independent simulations for each case were produced containing 10^6^ time steps of 0.5 ns at a temperature of 60*^◦^*C. Initially, particles were randomly distributed in a spherical volume of radius 200 nm. Interaction parameters were identical to the ones discussed earlier.

#### Detection and evaluation of simulated droplets

Droplets in the simulations were detected by using the DB scan algorithm as implemented in the python package scikit-learn (version 1.1.2). If not specified otherwise, a minimum sample number of 10 and a maximum distance of 40 nm between particles were used.

As a proxy for the volume of each droplet, the number of particles per droplet was calculated. Alternatively, we discretized the simulation box into cubes of 10 nm edge length and counted the number of filled boxes for each droplet. This approach introduces a limited resolution, as one voxel can contain multiple particles and structure is not resolved for scales smaller than the box length.

Sphericity was calculated by using the python package jakteristics in version 0.5.0 (see https://github.com/jakarto3d/jakteristics) which calculates the sphericity of a point cloud as ratio between the largest and the smallest eigenvector.

### Lattice model

The dispersal of Y-motif droplets by an addition of amphiphile-motifs was also studied by numerical simulation in a square lattice system with specific particle-particle interaction and excluded volume effects. The dynamic Monte-Carlo simulation is performed on a 64 *×* 64 two-dimensional lattice with periodic boundary conditions. Y-motifs are represented by T-shaped particles which occupy four sites in the lattice while amphiphiles and controls are denoted by cross shaped particles which occupy five lattice sites. The negative arm (red) of the amphiphiles is replaced by neutral arm (arm with a white dot in Figure 5E) for controls. We start from a random initial configuration with T-shaped particle concentration of 0.020 and amphiphilic concentration of 0.030. Due to the arrest effect of the particles resulting from their shape, a higher concentration of cross shaped motifs compared to the T-shaped motifs is needed to observe the amphiphilic effect in the system. The attractive interaction between any two arms except the negative arm is given by 3.0*k_B_T* while the repulsive interaction between the negative arm and other is 6.5*k_B_T,* where *T* is the explicit system temperature and the Boltzmann constant (*k_B_*) is taken as unity. The neutral arm is non-interacting, it only occupies a lattice site. A single Monte Carlo sweep (MCS) is defined when on average each DNA motif is attempted to move to any of its four directions with equal probability representing translational diffusion and attempted to rotate by *π/*4 angle in either direction with equal probability as rotational diffusion. We have simulated the system for maximum of 10^8^ MCS. All the movements and rotations took place using standard Metropolis algorithm, which involved change in interaction energy Δ*E* and the temperature *T* of the system. We have kept *T* = 0.45 fixed while increasing *T* would more likely randomize the system by thermal moves. We have measured area and circularity of the droplets from the snapshots of the system configuration using connected components function in MATLAB. To compare the results with experiment and with the Brownian dynamics simulation, we measured circularity which is analogous to sphericity in 3-dimensional system. The number of DNA motifs present in a given droplet was directly obtained via the droplet area in the unit of pixels, with each pixel representing one lattice site.

---

**Figure S1:**
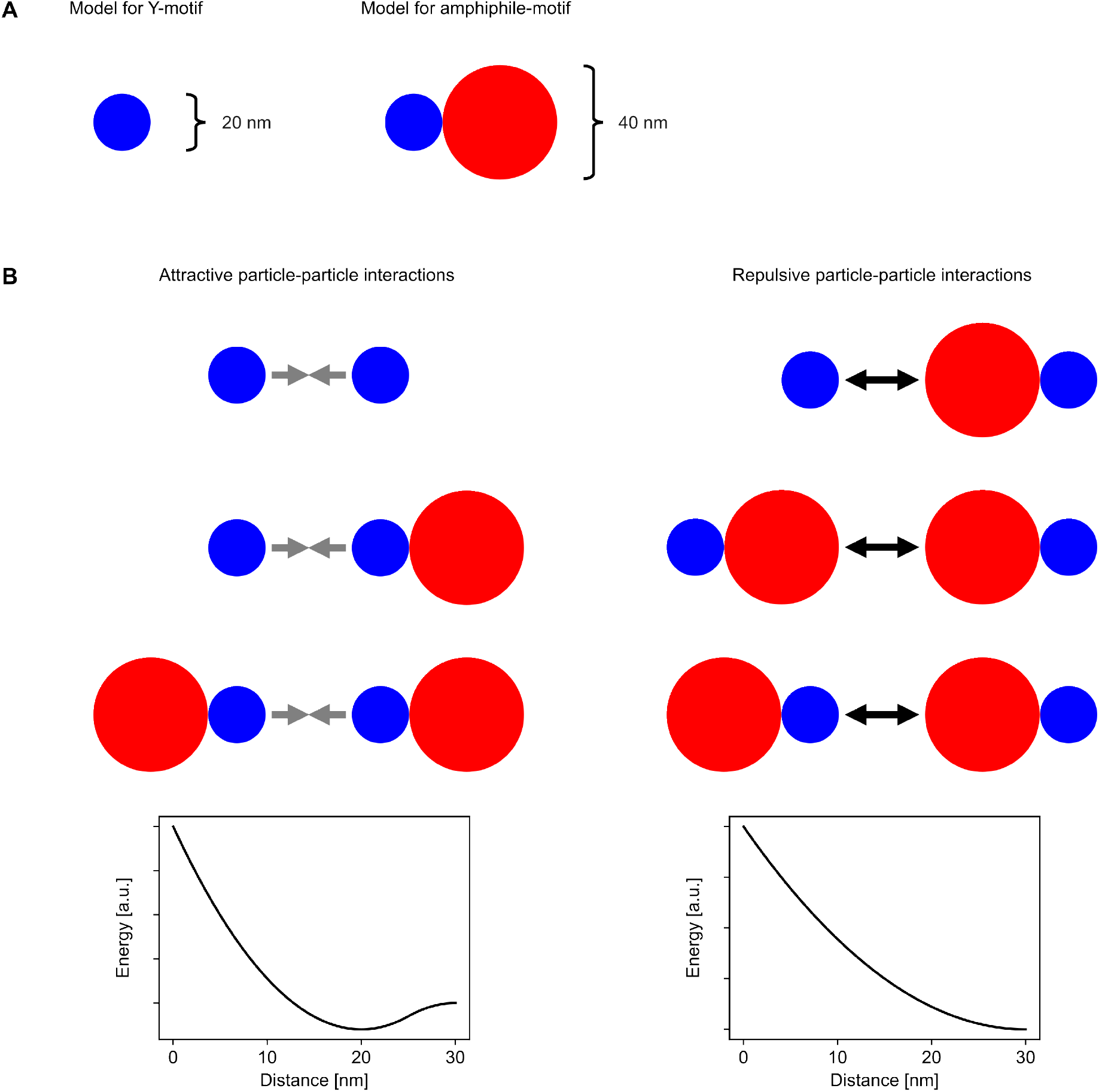
Summary of sizes and interaction parameters used for interacting-particle simulations. A) To-scale drawing of the two types of particles used in the simulation. B) Overview of attractive and repulsive particle interactions, displayed with interaction potentials that implement these interactions in the ReaDDy framework. For better visualisation, the energy axis uses arbitrary units and is not drawn to scale.

**Table S1:**
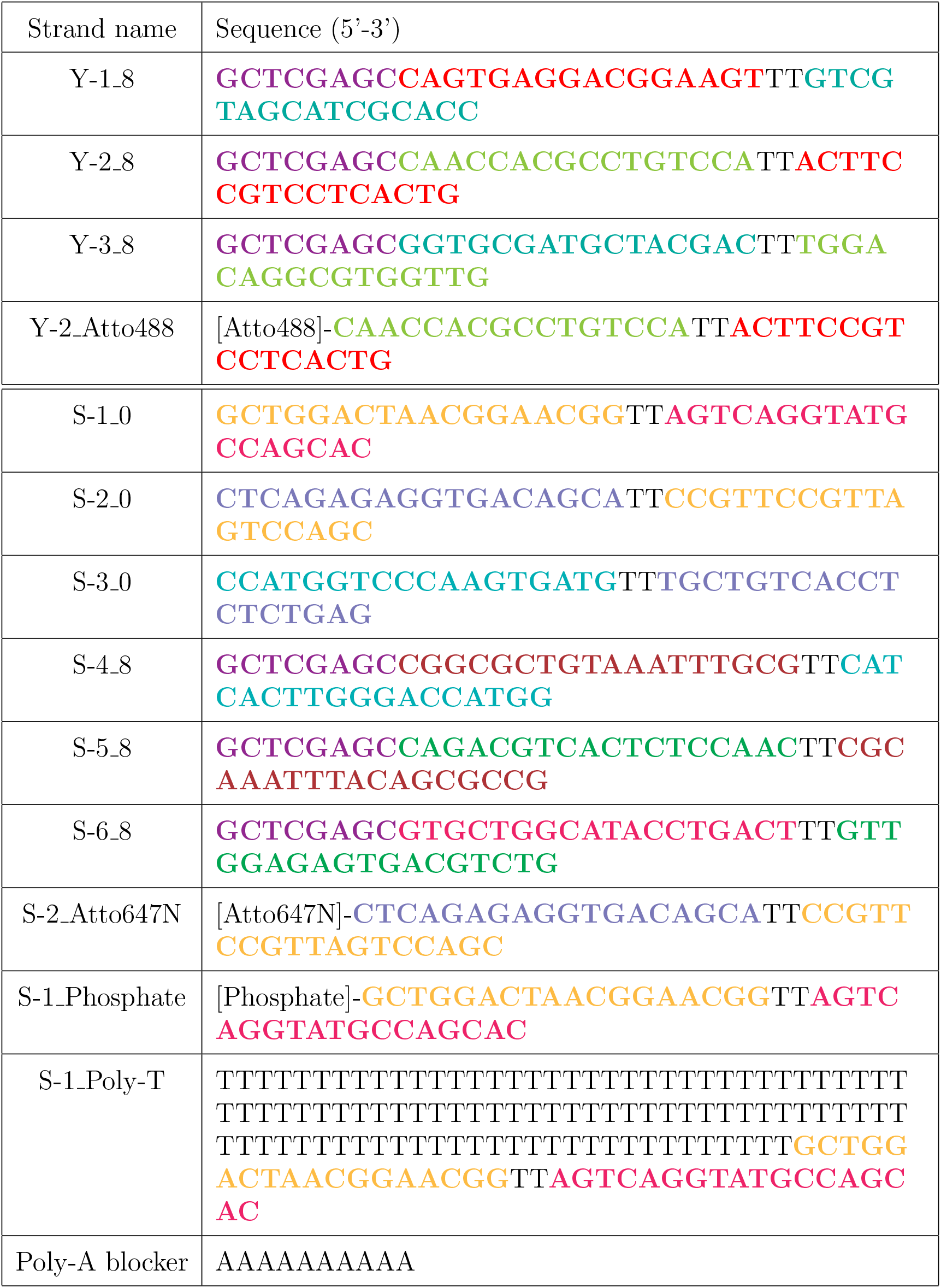
Strand names and sequences of nanomotifs

**Figure S2:**
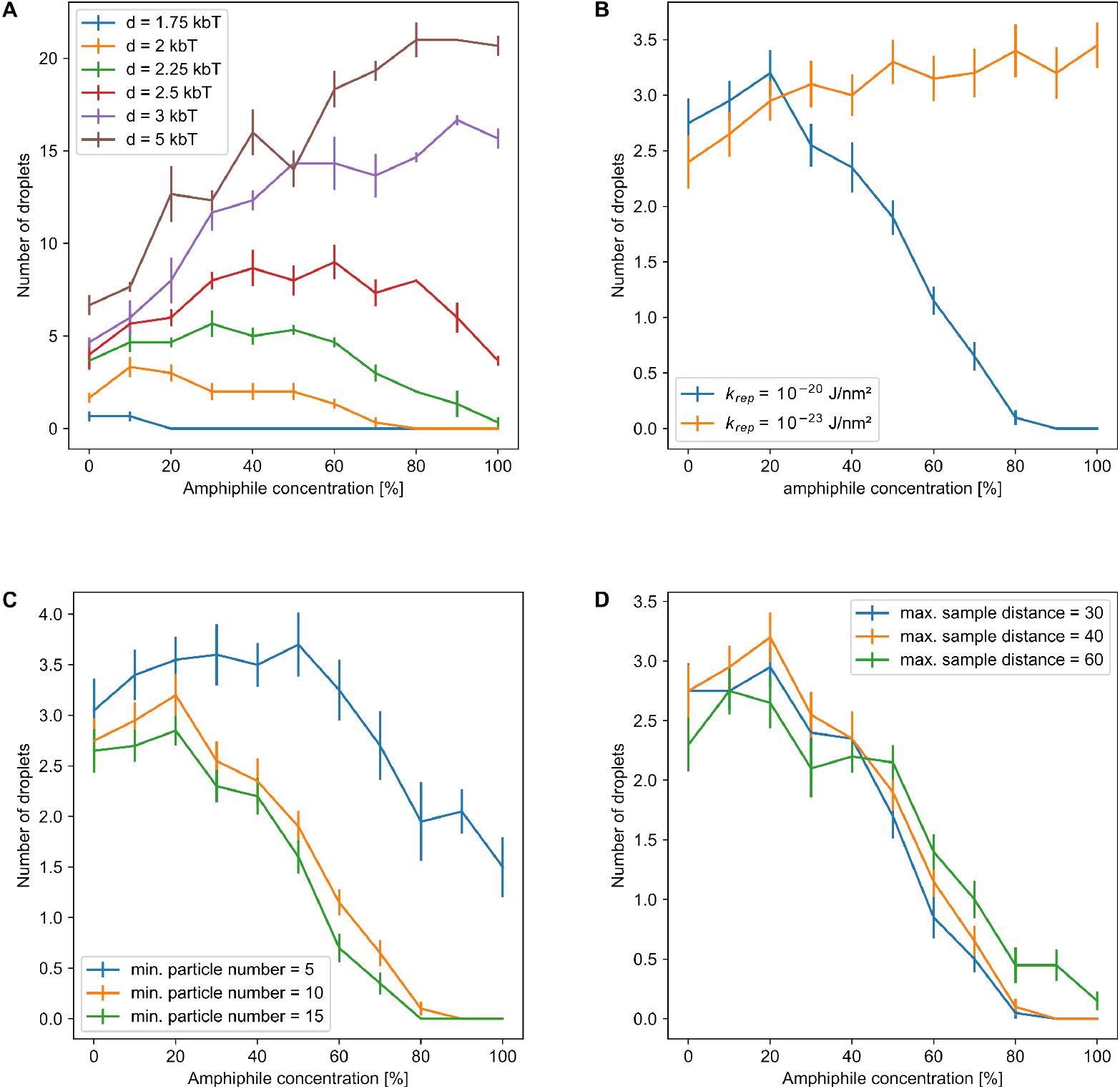
The number of detected droplets in the Brownian dynamics simulation for different simulation and evaluation parameters. A) To select the parameters of the Brownian dynamics simulation, we carried out out a simulated titration experiment that places 450 particles split between Y-motif and amphiphile in a spherical volume of radius 200 nm inside a simulation box of 1 *µm* edge length. We then examined the final number of droplets after 10^6^ time steps. As the depth of the harmonic well potential introducing attraction between simulated Y-motifs increases, the peak number of detected droplets shifts towards higher amphiphile concentrations. We selected a value of 2*k_b_T* as the resulting curve resembles the experimental observation. B) Keeping the depth of the well potential fixed at 2*k_b_T* and decreasing the repulsion between ampihpihle and all other particles resulted in a continuous increase of droplet number, confirming that the ration of attractive to repulsive interactions determines the shape of the curve. C) Droplets were detected with the DB-scan algorithm using a minimum number of 10 particles for classification as cluster/droplet. Changing this value by 5 particles influences the shape of the curve but not the qualitative behaviour. D) In a similar fashion, the maximum distance between particles of one cluster can be varied from the 40 nm we settled on, without changing the qualitative behaviour of the curve. Panel A averages over 3 independent simulation runs, panels B, C, and D use 20 simulation runs with well potential depth of 2*k_b_T* .

**Figure S3:**
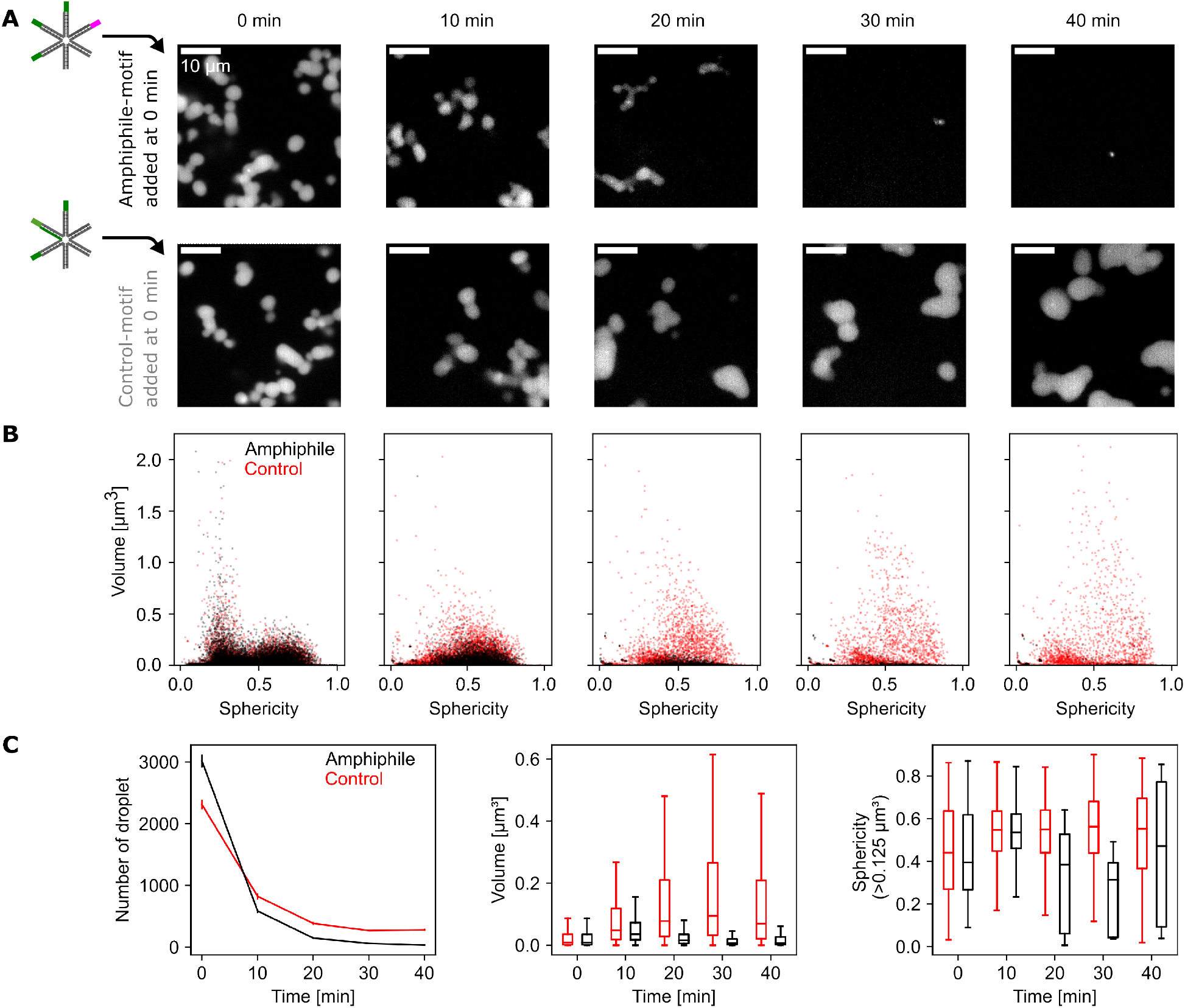
Time-course of droplet dispersal upon addition of amphiphile-motifs. A) Representative confocal microscopy sections showing the distribution of Y-motifs in flow cells prepared at 10-min time intervals after amphiphile-motifs were added to preformed Y-motif droplets (0 min). Several tubes prepared in this manner were maintained at a stable temperature of 50*^◦^*C, control-motif samples were processed alongside amphiphile-motifs experiments. B) Sphericity-volume distributions obtained from the analysis of confocal images. Three-dimensional analysis was carried out in image stacks (image stacks were acquired at 7 fields of view per sample and time point, data pooled from one experiment. Number of droplets, amphiphile-motif addition: n=21123, 4110, 1044, 437, 239; control-motif addition: n=6191, 5775, 2707, 1890, 1969. C) Time courses of the number of droplets per field of view (mean±SEM, *n* = 7 fields of view per time point), volume of droplets (standard boxplots, all droplet volumes included), and the sphericity of large droplets (volumes ≥ 0.125 *µ*m^3^ included).

**Figure S4:**
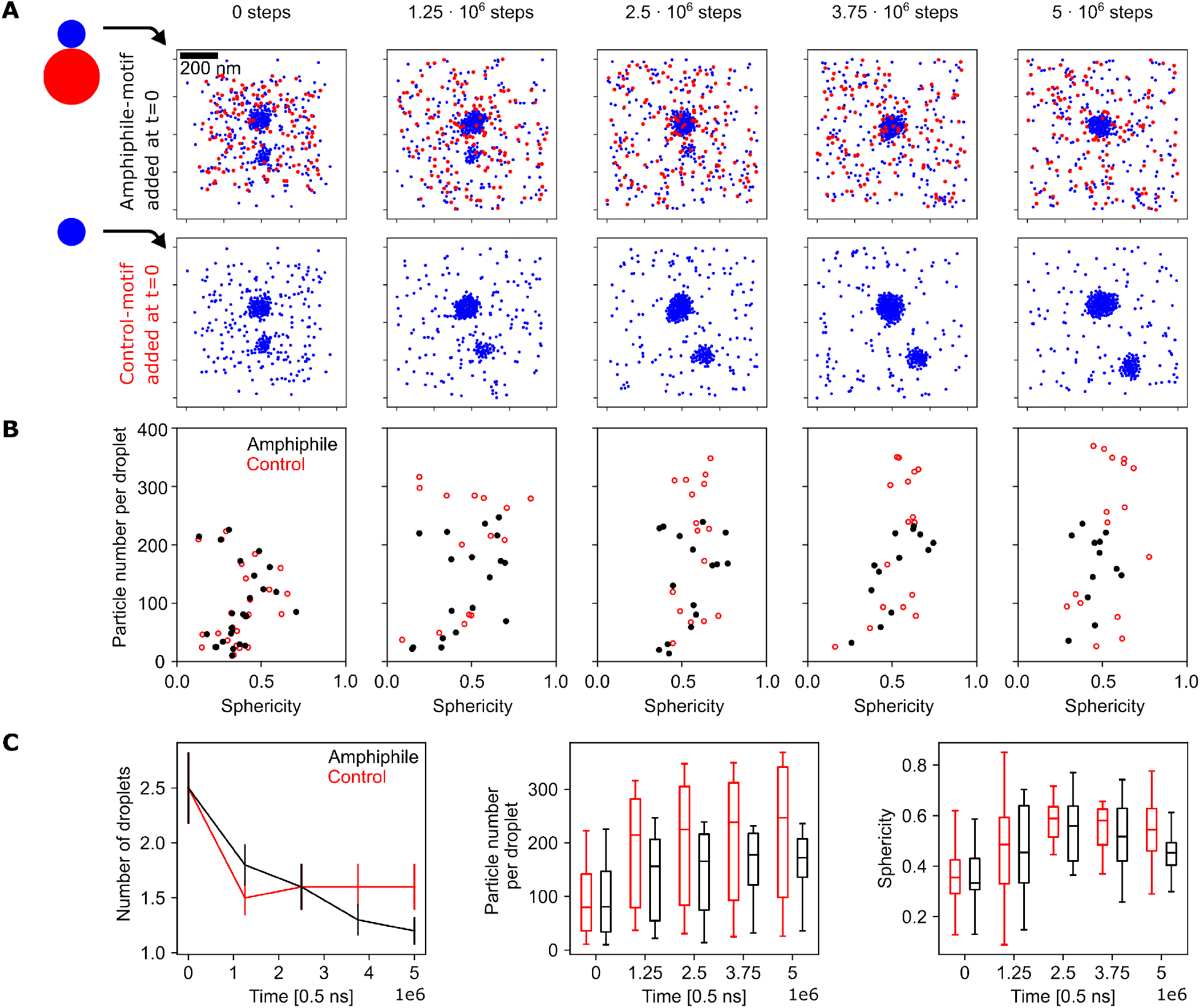
Time series of amphiphile-mediated droplet dispersal obtained from interacting-particle simulations. A) To simulate the time-lapse experiment, either amphiphile-motifs or control-motifs were added to a simulation containing preformed Y-motif droplets. The preformed droplets were generated by adding 450 Y-motifs to a simulation box of 1x1x1 *µ*m and simulating for 10^6^ steps at a temperature of 60*^◦^*C. Following this step, 450 additional amphiphiles or Y-motifs serving as control were added at random positions in a way that particle overlaps are prevented. 5 10^6^ additional steps at a temperature of 40*^◦^*C were then carried out. Representative z-slices of 200 nm thickness from both simulation cases are shown. Both simulations started from the same initial configuration. Control motifs were implemented identically to Y-motifs and are also displayed in blue color. B) 10 simulations for each case were used to generate distributions of sphericity versus particle number per droplet. C) The number of droplets per simulation (mean±SEM, *n* = 10 simulations) and boxplots for particle number per droplet and sphericity are shown for different time points.

**Figure S5:**
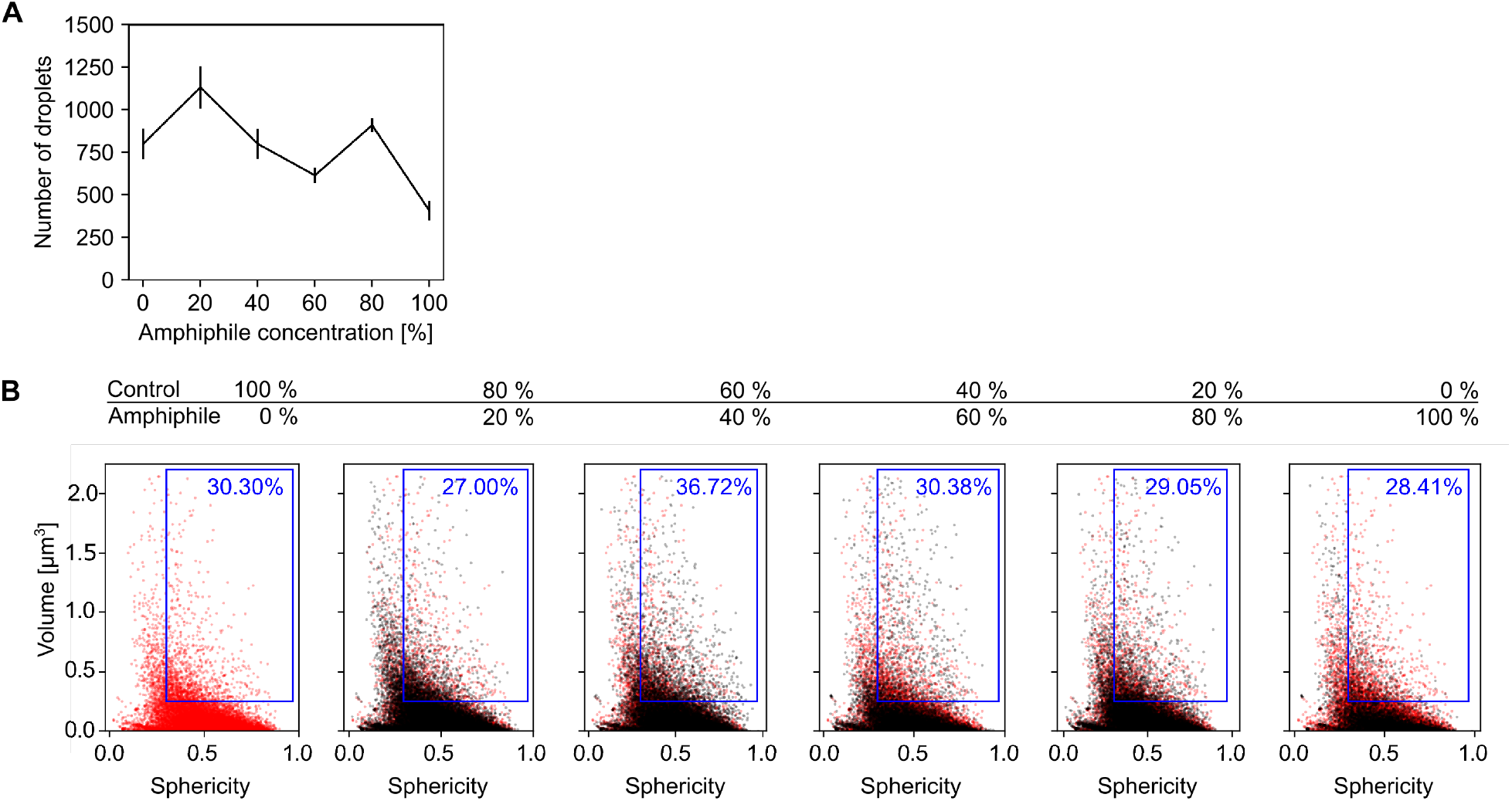
Repeat of the amphiphile titration experiment. A repeat of the titration experiment described in figure 4 resulted in similar behaviour.Y-motifs, control motifs (no poly-T extension), and amphiphile-motifs were separately prepared, mixed at different percentages of amphiphile motif, transiently heated (60*^◦^*C), and injected into flow cells for microscopy (ambient temperature 40*^◦^*C). As the volume fraction of amphiphile increases, the number of droplets and the percentage of larger droplets with high sphericity transiently increases. A) Changes of the number of droplets per field of view with amphiphile concentration (mean SEM, *n* = 14, 13, 14, 14, 14, 14 fields of view). B) Sphericity-volume distributions obtained from the analysis of confocal image stacks (image stacks were acquired at up to five fields of view per sample, n=11206, 14718, 11207, 8609, 12740, 5708 droplets, data pooled from *N* = 14, 14, 13, 13, 14, 14 total fields of view pooled from three independent experimental repeats). Small droplets (volume≤ 0.125 *µ*m^3^) were excluded in calculation of percentages.

**Figure S6:**
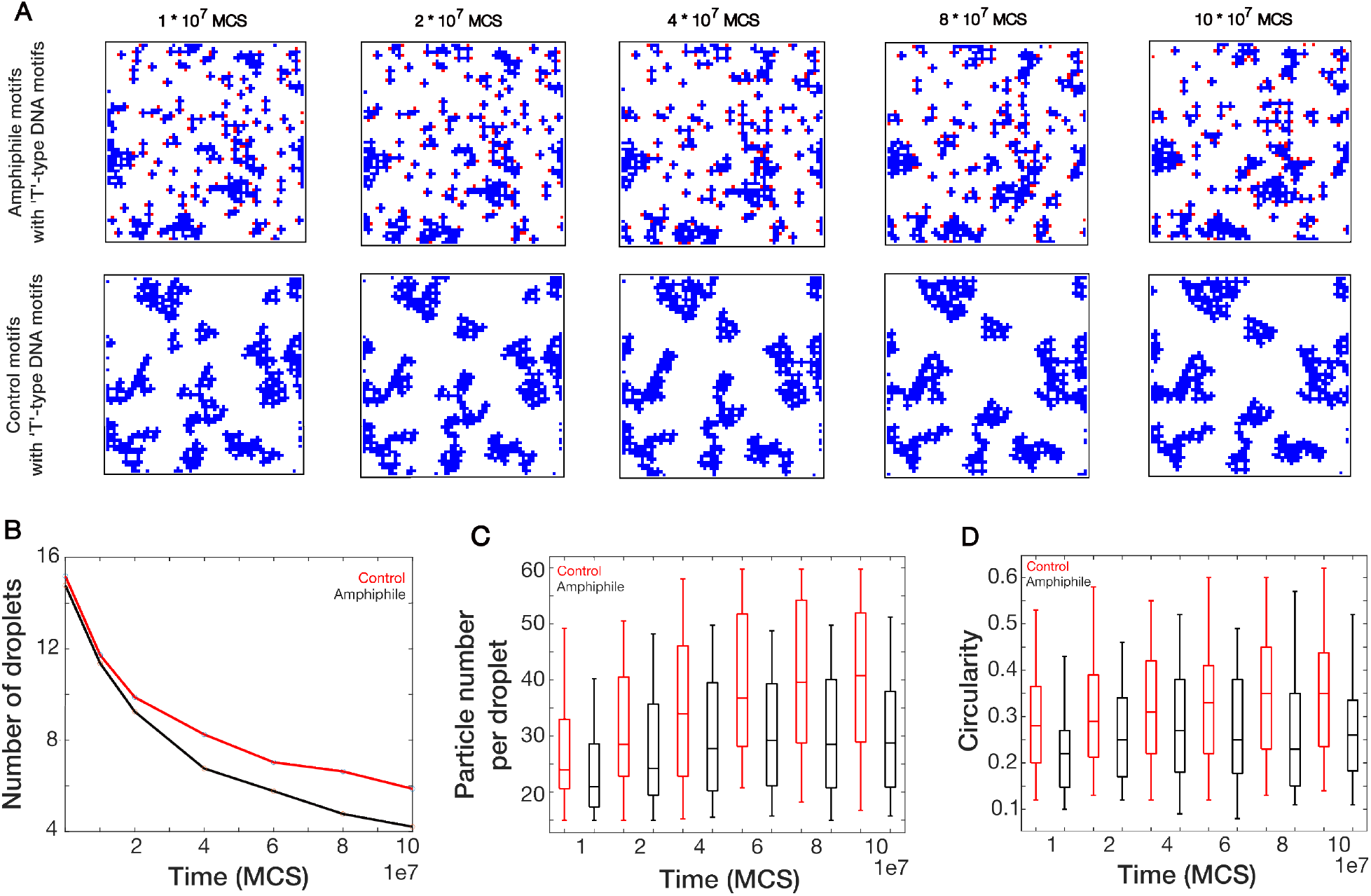
Analysis of lattice simulation. A) Time-lapse snapshots for lattice simulation are illustrated when amphiphile and control motifs are introduced in the system of T-shaped DNA motifs separately. The simulations has been carried out on a 64 *X* 64 lattice system. B) Mean number of droplets per simulation (mean±SEM, *n* = 25 simulations) is shown as time progresses and C,D) Boxplots for time-lapse simulation are presented for particle number per droplet and circularity respectively.

