## Supplementary material for "Amphiphiles formed from synthetic DNA-nanomotifs mimic the step-wise dispersal of transcriptional clusters in the cell nucleus": OligomerSequences

**DNA-nanomotif sequences**

**Y-1_8**

5’- GCT CGA GCC AGT GAG GAC GGA AGT TTG TCG TAG CAT CGC ACC -3’

**Y-2_8**

5’- GCT CGA GCC AAC CAC GCC TGT CCA TTA CTT CCG TCC TCA CTG -3’

**Y-3_8**

5’- GCT CGA GCG GTG CGA TGC TAC GAC TTT GGA CAG GCG TGG TTG -3’

**Y-2_0_Atto488**

5'- Atto488 - CAA CCA CGC CTG TCC ATT ACT TCC GTC CTC ACT G -3'

**orthY-1_8**

5’- CTC GCG AGA AAG GAA CTC TCC GCG TTG ACA AAG CCG ACA CGT -3’

**orthY-2_8**

5’- CTC GCG AGG CCT CTG TGT CGC ATC TTC GCG GAG AGT TCC TTT -3’

**orthY-3_8**

5’- CTC GCG AGA CGT GTC GGC TTT GTC TTG ATG CGA CAC AGA GGC -3’

**orthY-2_0_Atto594**

5' Atto594-GCC TCT GTG TCG CAT CTT CGC GGA GAG TTC CTT T -3'

**S-1_8**

5’- CTC GCG AGG CTG GAC TAA CGG AAC GGT TAG TCA GGT ATG CCA GCA C -3’

**S-2_8**

5’- CTC GCG AGC TCA GAG AGG TGA CAG CAT TCC GTT CCG TTA GTC CAG C -3’

**S-3_8**

5’- CTC GCG AGC CAT GGT CCC AAG TGA TGT TTG CTG TCA CCT CTC TGA G -3’

**S-4_8**

5’- GCT CGA GCC GGC GCT GTA AAT TTG CGT TCA TCA CTT GGG ACC ATG G -3’

**S-5_8**

5’- GCT CGA GCC AGA CGT CAC TCT CCA ACT TCG CAA ATT TAC AGC GCC G -3’

**S-6_8**

5’- GCT CGA GCG TGC TGG CAT ACC TGA CTT TGT TGG AGA GTG ACG TCT G -3’

**S-2_Atto647N**

5’- Atto647N - CTC AGA GAG GTG ACA GCA TTC CGT TCC GTT AGT CCA GC -3’

**S-1_0**

5’- GCT GGA CTA ACG GAA CGG TTA GTC AGG TAT GCC AGC AC -3’

**S-2_0**

5’- CTC AGA GAG GTG ACA GCA TTC CGT TCC GTT AGT CCA GC -3’

**S-3_0**

5’- CCA TGG TCC CAA GTG ATG TTT GCT GTC ACC TCT CTG AG -3’

**S-1_Poly-T**

5’- TTT TTT TTT TTT TTT TTT TTT TTT TTT TTT TTT TTT TTT TTT TTT TTT TTT TTT TTT TTT TTT TTT TTT TTT TTT TTT TTT TTT TTT TTT TTT TTT TTT TTT GCT GGA CTA ACG GAA CGG TTA GTC AGG TAT GCC AGC AC -3’

**S-1_Phosphate**

5’- Phosphate-GCT GGA CTA ACG GAA CGG TTA GTC AGG TAT GCC AGC AC -3’

**Poly-A blocker**

5’- AAA AAA AAA A -3’

Tab. 1: Strand names and sequences of nanomotifs

| Strand name | Sequence (5’-3’) |
| --- | --- |
| Y-1 8 | GCTCGAGCCAGTGAGGACGGAAGTTTGTCG |
|  | TAGCATCGCACC |
| Y-2 8 | GCTCGAGCCAACCACGCCTGTCCATTACTTC |
|  | CGTCCTCACTG |
| Y-3 8 | GCTCGAGCGGTGCGATGCTACGACTTTGGA |
|  | CAGGCGTGGTTG |
| Y-2 Atto488 | [Atto488]-CAACCACGCCTGTCCATTACTTCCGT |
|  | CCTCACTG |
| S-1 0 | GCTGGACTAACGGAACGGTTAGTCAGGTATG |
|  | CCAGCAC |
| S-2 0 | CTCAGAGAGGTGACAGCATTCCGTTCCGTTA |
|  | GTCCAGC |
| S-3 0 | CCATGGTCCCAAGTGATGTTTGCTGTCACCT |
|  | CTCTGAG |
| S-4 8 | GCTCGAGCCGGCGCTGTAAATTTGCGTTCAT |
|  | CACTTGGGACCATGG |
| S-5 8 | GCTCGAGCCAGACGTCACTCTCCAACTTCGC |
|  | AAATTTACAGCGCCG |
| S-6 8 | GCTCGAGCGTGCTGGCATACCTGACTTTGTT |
|  | GGAGAGTGACGTCTG |
| S-2 Atto647N | [Atto647N]-CTCAGAGAGGTGACAGCATTCCGTT |
|  | CCGTTAGTCCAGC |
| S-1 Phosphate | [Phosphate]-GCTGGACTAACGGAACGGTTAGTC |
|  | AGGTATGCCAGCAC |
| S-1 Poly-T | TTTTTTTTTTTTTTTTTTTTTTTTTTTTTTTTTTTT |
|  | TTTTTTTTTTTTTTTTTTTTTTTTTTTTTTTTTTTT |
|  | TTTTTTTTTTTTTTTTTTTTTTTTTTTTTTGCTGG |
|  | ACTAACGGAACGGTTAGTCAGGTATGCCAGC |
|  | AC |
| Poly-A blocker | AAAAAAAAAA |
